# Neural stem cells induce the formation of their physical niche during organogenesis

**DOI:** 10.1101/149955

**Authors:** Ali Seleit, Isabel Krämer, Bea Riebesehl, Elizabeth M. Ambrosio, Julian S. Stolper, Nicolas Dross, Colin Q. Lischik, Lazaro Centanin

## Abstract

Most organs rely on stem cells to maintain homeostasis during post-embryonic life. Typically, stem cells of independent lineages work coordinately within mature organs to ensure proper ratios of cell types. Little is known, however, on how these different stem cells locate to forming organs during development. Here we show that neuromasts of the posterior lateral line in medaka are composed of two independent life-long lineages with different embryonic origins. Clonal analysis and 4D imaging revealed a hierarchical organisation with instructing and responding roles: an inner, neural lineage induces the formation of an outer, border cell lineage (nBC) from the skin epithelium. Our results demonstrate that the neural lineage is necessary and sufficient to generate nBCs highlighting self-organisation principles at the level of the entire embryo. We hypothesise that transformation of surrounding tissues plays a major role during the establishment of vertebrate stem cell niches.

## Introduction

Animal organs are composed of diverse cell types, which in most cases derive from different embryonic origins. The last decades have witnessed huge efforts focused on identifying molecules necessary for organ formation in a wide range of organisms. This approach resulted in a deeper understanding of the molecular networks that control organogenesis in both vertebrates and invertebrates (Gilbert, 2014). Even though a broad range of molecular details have been uncovered, those did not contribute to revealing how cells from different embryonic origins end up together in the same functional unit.

There are two alternative scenarios on how cells from different lineages successfully come together during embryogenesis to assemble composite organs. One option is that they originate from cells that form in remote areas and migrate to meet at a defined location. This typically happens during the formation of gonads in vertebrates, where germ cells follow a directional migration towards the somatic cells that will form their stem cell niche and eventually the gonad (Doitsidou *et al.*, 2002; Santos and Lehmann, 2004). The other option is that one cell type is generated in a stereotypic position to then induce the recruitment and transformation of a second cell type to fulfil a new role *in situ.* This occurs during eye formation, where retinal progenitors induce the transformation of surface ectodermal cells into lens cells that are later integrated into the forming eye (Gilbert, 2014).

While gonads, eyes and most animal organs have a defined number and location within species, the neuromasts - sensory organs of the lateral line system in fish - constitute a more plastic model in which organ numbers cannot exactly be deduced from the developmental stage or overall size (Ghysen and Dambly-Chaudière, 2007; Seleit *et al.*, 2017). The first neuromasts of the posterior lateral line (pLL) system are generated during embryogenesis by one primordium (in medaka) and two primordia (in zebrafish, tuna and *Astyanax*) that migrate anterior-to-posterior along the horizontal myoseptum while depositing groups of cells from their trailing ends at regular intervals. Soon after deposition these cellular clusters mature into neuromasts, composed of three main cell types as assessed by morphology and molecular markers. Hair cells (HCs), the sensory component and differentiated cell type, project cilia into the external environment and are responsible for detecting water flow and relaying this information back to the CNS (Ghysen and Dambly-Chaudière, 2007; Williams and Holder, 2000). Surrounding the HCs is a population of progenitor support cells (SCs) (Ghysen and Dambly-Chaudière, 2007; Hernández *et al.*, 2007), while mantle cells (MCs) form an outer ring around both cell types and encapsulate the projecting HCs in a cupula-like structure (Jones and Corwin, 1993; Steiner *et al.*, 2014). Since HCs are continuously lost and replaced under homeostatic conditions (Cruz *et al.*, 2015; Williams and Holder, 2000) and in response to injury (Hernández *et al.*, 2007; López-Schier and Hudspeth, 2006) throughout the life of fish (Pinto-Teixeira *et al.*, 2015), the existence of stem cells capable of constantly replenishing HCs has been proposed.

The presence of neuromast stem cells has also been inferred from the fact that new neuromasts are continuously added to the lateral line system during the lifetime of fish, presumably to cope with an increasing body size (Dufourcq *et al.*, 2006; Ghysen and Dambly-Chaudière, 2007; Wada *et al.*, 2013). It has been reported that the peripheral ring of mantle cells plays a major role in new organ formation (Dufourcq *et al.*, 2006; Jones and Corwin, 1993; Stone, 1933; Wada *et al.*, 2013). This has led to the proposition of a linear model in which mantle cells give rise to support cells that in turn replenish lost hair cells (Ghysen and Dambly-Chaudière, 2007). The transition from support to hair cells has been heavily investigated in the past decade and is strongly supported by a large body of experimental evidence (Hernández *et al.*, 2007; López-Schier and Hudspeth, 2006; Ma *et al.*, 2008; Romero-Carvajal *et al.*, 2015; Wibowo *et al.*, 2011; Williams and Holder, 2000). However, the proposed transition of mantle into support cells has not yet been shown and it is still unknown where the stem cells of the neuromasts reside (Pinto-Teixeira *et al.*, 2015). An interesting observation is that upon repeated injury of the hair cell population coupled with a depletion of support cells, mantle cells re-enter the cell cycle (Romero-Carvajal *et al.*, 2015), but their precise role during homeostasis and regeneration remains unclear. The current view posits support cells as multipotent progenitors capable of self-renewing and differentiation and suggests a niche role for mantle cells influencing SC proliferative behaviour and fate (Romero-Carvajal *et al.*, 2015). In the absence of long-term lineage tracing data, however, it is very difficult to reveal the identity and location of neuromast stem cells and to assess if homeostatic replacement, response to injuries, and generation of new organs during post-embryonic stages are all performed by the same or different cell types.

Here we use newly developed transgenic lines to follow lineages during development and into adulthood in neuromasts of medaka fish. We prove that mantle cells constitute *bona fide* neuromast neural stem cells during homeostasis, growth and organ regeneration. Additionally, we identify a new population of neuromast cells that we name neuromast border cells (nBCs), which constitute a different lineage that never crosses boundaries with the neural lineage maintained by mantle cells. We track border cells back to earlier developmental stages and demonstrate that they do not originate from the pLL primordium but rather from the skin epithelium, defining neuromasts as composite organs. Finally, we show that neural stem cells are necessary and sufficient to induce the transformation of epithelial cells into nBCs, a process that we followed by genetic lineage analysis and 4D imaging. Altogether, we uncover that neural stem cells recruit accessory cells that will be maintained as a life-long separate lineage.

## Results

### nBCs are the outer cells of the organ

To address the existence and identity of neuromast stem cells we decided to follow a lineage analysis approach using the *Gaudí* toolkit (Centanin *et al.*, 2014), in combination with transgenic lines that label the different cell types within mature neuromasts. Tg(*Eya1*:EGFP) (Seleit *et al.*, 2017) is expressed in all cells of the migrating primordium and during organogenesis, but is restricted to hair cells and internal support cells located underneath HCs in mature organs (Figure 1A, A’). The newly generated enhancer trap *neurom^K8^*:H2B-EGFP stably labels all support cells and mantle cells (Figure 1B, B’)(See Materials & Methods). Additionally, the Tg(*K15*:H2B-RFP) labels a subset of the *neurom^K8^* positive cells that form a peripheral ring in mature neuromasts (Figure 1C-C’) and are Sox2 positive (Figure 1D-D”). A 3D reconstitution of triple transgenic Tg(*K15*:mEYFP), Tg(*K15*:H2B-RFP), Tg(*Eya1*:EGFP) neuromasts indicates that EYFP positive cells are wrapping the hair cells of the organ (Figure 1E and Supplementary Movie 1), a distinctive morphology and position that characterises mantle cells (Jones and Corwin, 1993; Steiner *et al.*, 2014). Both *neurom^K8^*:H2B-EGFP and Tg(*K15*:H2B-RFP) also label skin epithelia on the entire body of juvenile and adult medaka fish. This combination of transgenic lines allows us to dynamically asses the cell content of neuromasts *in vivo* during embryonic, juvenile and adult organ growth.

**Figure 1.**
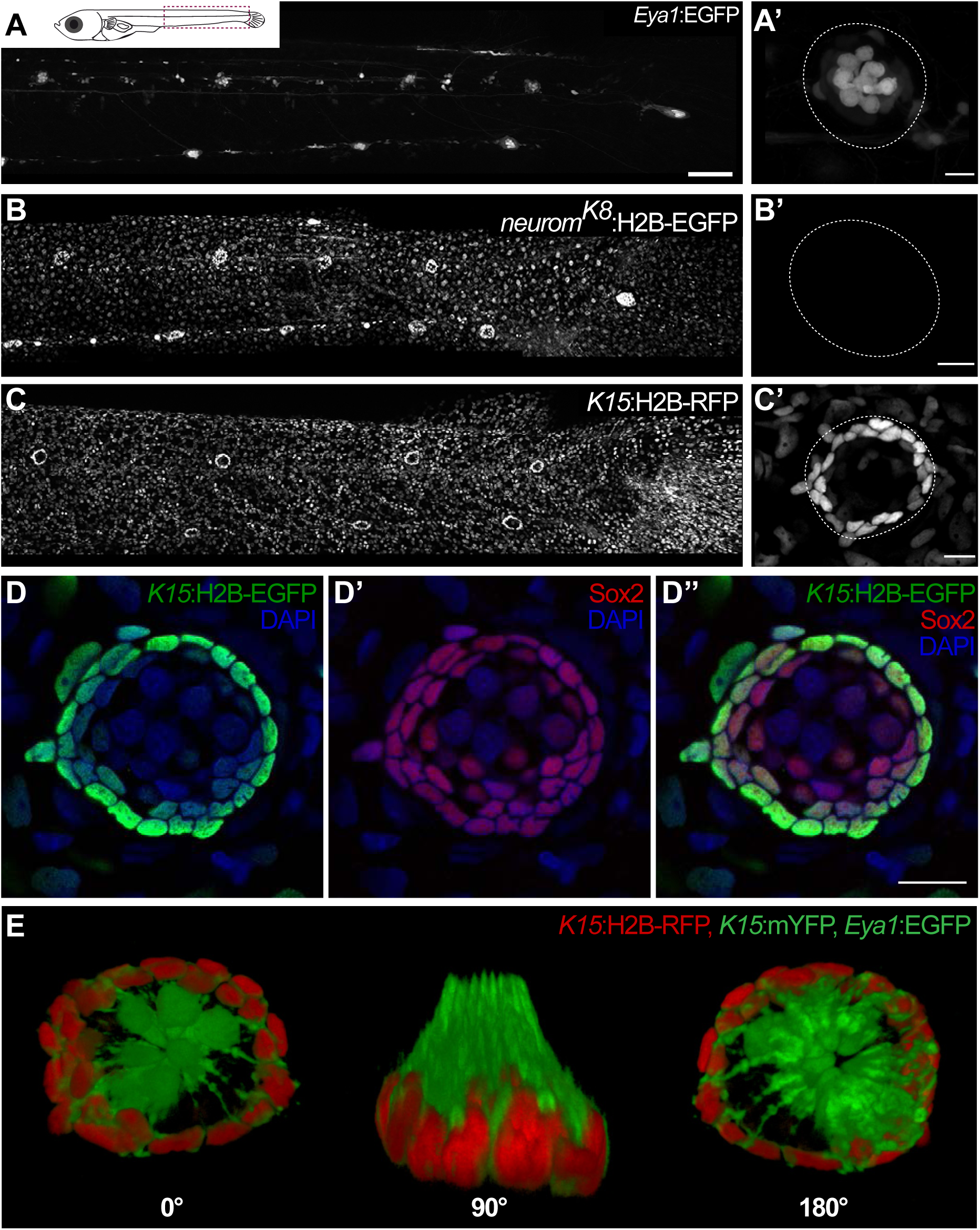
Specific transgenic lines label mantle, support and hair cells in mature medaka neuromasts. Tg(*Eya1*:EGFP) allows visualisation of all neuromasts along the pLL (A), and labels hair cells and internal support cells (A’). The enhancer trap *neuront^K8^* line labels skin epithelia (B) and mantle and support cells of a mature neuromast (B’). Tg(*K15*:H2B-RFP) also labels skin epithelia all over the body surface (C), but RFP expression in a mature neuromast is restricted to mantle cells (C’). *K15*^+^ mantle cells are Sox2+ as revealed by immunostainings (D-D”). A 3D reconstruction of a mature neuromast (E) of the triple transgenic line Tg(*K15*:mYFP) Tg(*Eya1*:EGFP), Tg(*K15*:H2B-RFP) of an early juvenile. Six neuromast hair cells (green bundles) project outwards, surrounded by a ring of mantle cells (red nuclei and green membranes) that encapsulate the hair cell bundles in a cupula-like structure. Scalebars are 100*μ*m for entire trunks (A, B, C) and 10*μ*m in neuromast close-ups.

While observing neuromasts counterstained with DAPI, we noticed that *K15* positive (*K15^+^)* mantle cells are consistently surrounded by an outer ring of elongated nuclei (Figure 2A, A’). This is the case for all neuromasts in medaka, including ventral, midline and dorsal neuromasts on the posterior lateral line, and neuromasts of the anterior lateral lines in both juveniles and adults (N > 100 neuromasts). Since these elongated nuclei locate to the outer border of neuromasts, we named the corresponding cells neuromast Border Cells (nBCs). Electron microscopy revealed that the membranes of border cells are intimately associated with those of mantle cells, often producing cytoplasmic protrusions into one another (Figure 2B-B”’) In addition we also observed desmosomes between MCs and nBCs (Figure C-C”). Using iterative imaging on Tg(*K15*:H2BEGFP) medaka larvae, we detected that a proportion of nBCs are EGFP positive (Figure 2D) but this expression decays within the following days. The transient nature of this expression suggests nBCs could originate from *K15^+^* cells and inherit the fluorescent protein, which in this case would be acting as a short-term lineage tracer. We therefore focused on revealing the embryonic origin and lineage relations of all neuromast cell types (Figure 2E) during homeostatic maintenance, organ growth and post-embryonic organogenesis.

**Figure 2.**
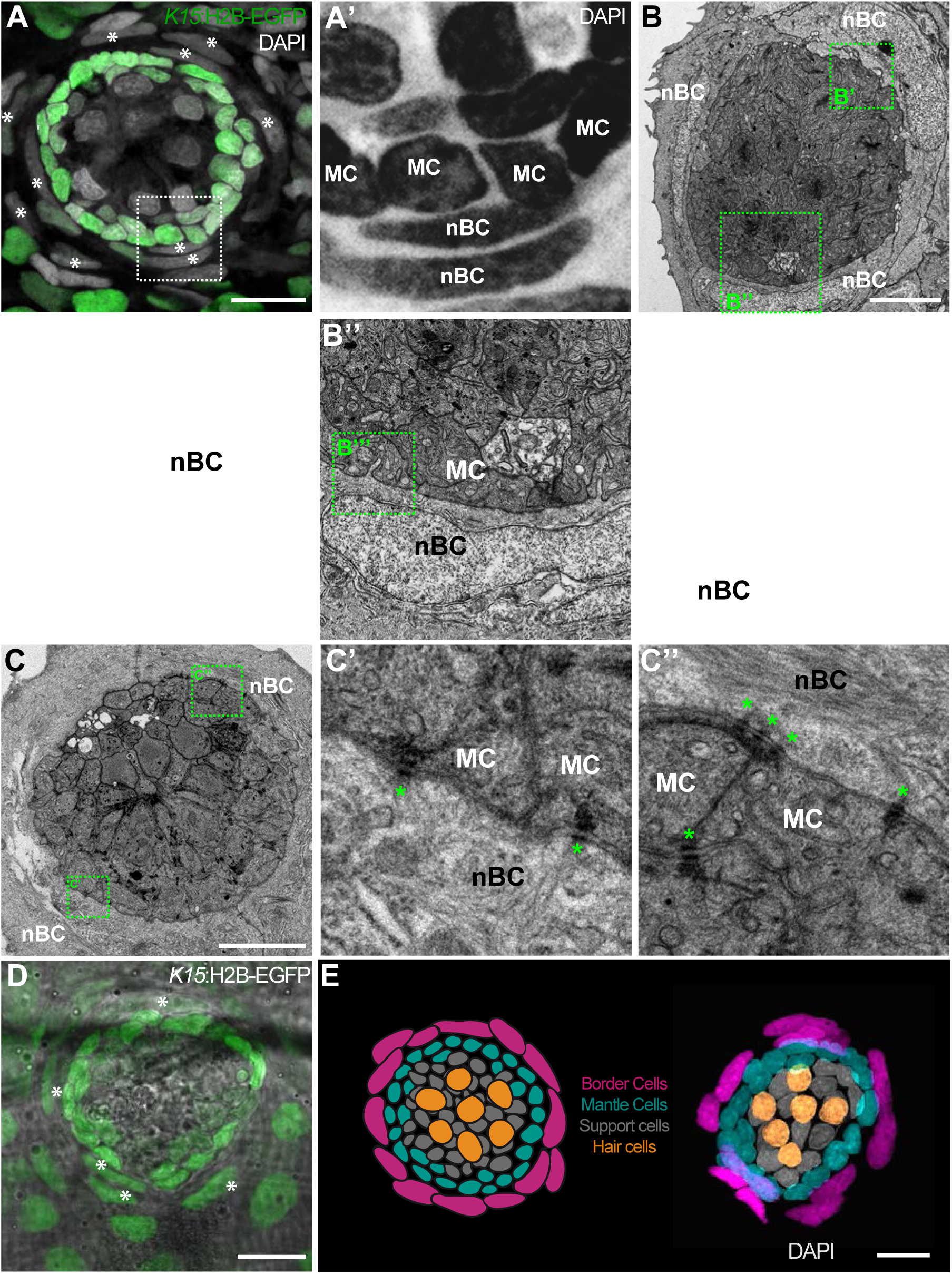
nBCs surround mantle cells of the neural lineage in mature neuromasts. An early juvenile neuromasts from Tg(*K15*:H2B-GFP) shows mantle cells (green in A, “MC” in A’) that are closely surrounded by elongated nuclei (border cells, nBCs) visualised by DAPI (white asterisks in A, “nBC” in A’). DAPI is shown in grey (A) and black (A’) to enhance contrast. Electronmicroscopy reveals that nBC and K15+ mantle cells are in close contact (B-C”). (B) Overview of a mature neuromast where mantle cells are surrounded by 3 nBCs. (B’-B”’) Zoom-in panels from figure (B) reveal a close association between MCs and nBCs that includes cytoplasmic protrusions of each cell type into the other (B’-B”’). (C) An upper section on a neuromast where mantle cells are surrounded by two nBCs. The darker dots (green asterisks) are cytoplasmic plaques of desmosomes formed between mantle cells and nBCs (C’, C”). (D) A younger neuromast than the one depicted in (A) from Tg(*K15*:H2B-GFP) shows that nBCs are also labelled with GFP (white asterisks in D). (E) *Pseudo-coloured* DAPI neuromast and scheme depicting the four cell types observed in every mature neuromast organ. Hair cells are shown in yellow, support cells in grey, mantle cells in green and border cells in magenta. Scalebars are 10*μ*m.

### nBCs constitute an independent life-long lineage

To understand the lineage relations between the different cells types of mature neuromasts, we decided to label individual cells in neuromasts and follow clones over time using the Gaudí toolkit (Centanin *et al.*, 2014). We generated positive clones by inducing recombination in *Gaudí^RSG^* (*ubiquitin*:LoxP-DsRed-LoxP-H2B-EGFP) crossed to either *Gaudí^Ubiq.iCre^* (*ubiquitin*:Cre*^ERT2^*) or *Gaudí^Hsp70.A^* (*Hsp70*:_nls_Cre) embryos and followed these for up to 19 days (Figure 3A). We imaged single clones in neuromasts of the posterior lateral line system two days post-induction, and then again 7 to 19 days after. The analysis of lineages of 144 clones in 91 neuromasts suggested two independent short-term lineages. One lineage involves mantle cells that expand and generate support cells and hair cells (Figure 3B-C”) - the neural lineage - (45,1%, 65/144 clones) while the other lineage was restricted to neuromast border cells (Figure 3D-D”) (12,5%, 18/144 clones).

**Figure 3.**
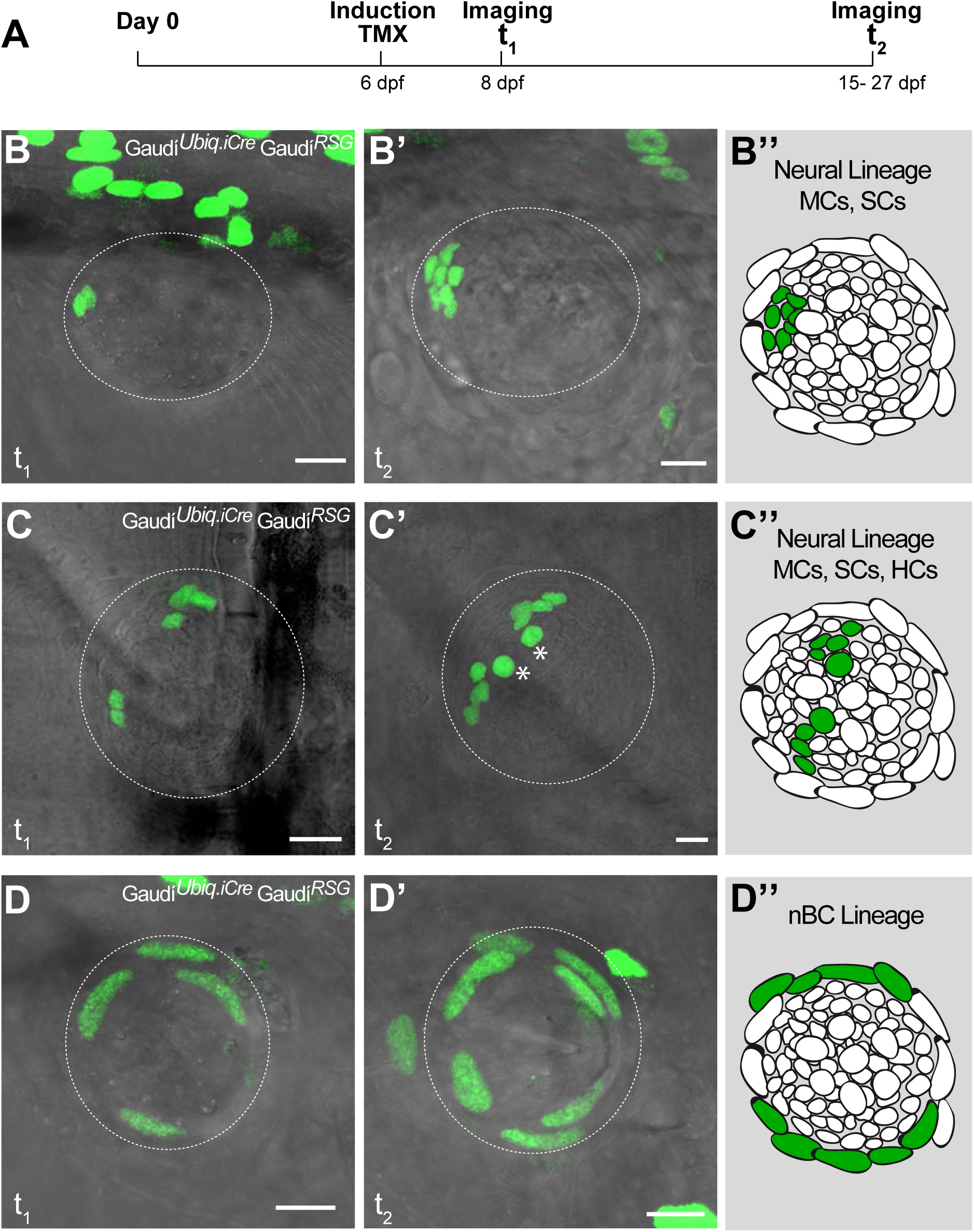
Mature neuromasts are composed of two separate lineages. (A) Experimental outline of clonal induction and short-term lineage tracing. t2 = 15 dpf. for B’ and D’, and 27 dpf for C’. (BD”) Iterative imaging of the same clones (H2B-EGFP) after recombination via tamoxifen reveals different behaviour of neuromast cells. Live imaging of clones containing mantle cells at t1, which expanded into the support cell domain (B,B’ and scheme in B”) and even further to generate hair cells (C, C’ and scheme in C”). Clones that contain labelled nBCs at t1 were restricted to the nBCs domain later on (D, D’ and scheme in D”), revealing a separate lineage for MCs-SCs-HCs and nBCs. Scalebars are 10*μ*m.

To explore whether this fate-restriction is maintained life-long, we focused on the *caudal neuromast cluster* (CNC) because of its stereotypic location on the caudal fin (Wada *et al.*, 2008). The CNC contains an increasing amount of neuromasts as fish age - the older the fish, the more neuromasts in the CNC. Neuromasts in the CNC are generated post-embryonically, presumably from a founder, embryonic neuromast - neuromast^P0^ (Figure 4A, B). We confirmed this by two photon laser ablations, which revealed that eliminating the neuromast^P0^ at eight days post-fertilisation (dpf)(Figure 4C, C’) results in an adult missing the CNC on the experimental side but with a wild-type CNC on the contralateral, control side (Figure 4D, E). The CNC represents a system that allows us to investigate neuromast stem cells during organ growth, homeostasis and also during post-embryonic organ formation.

**Figure 4.**
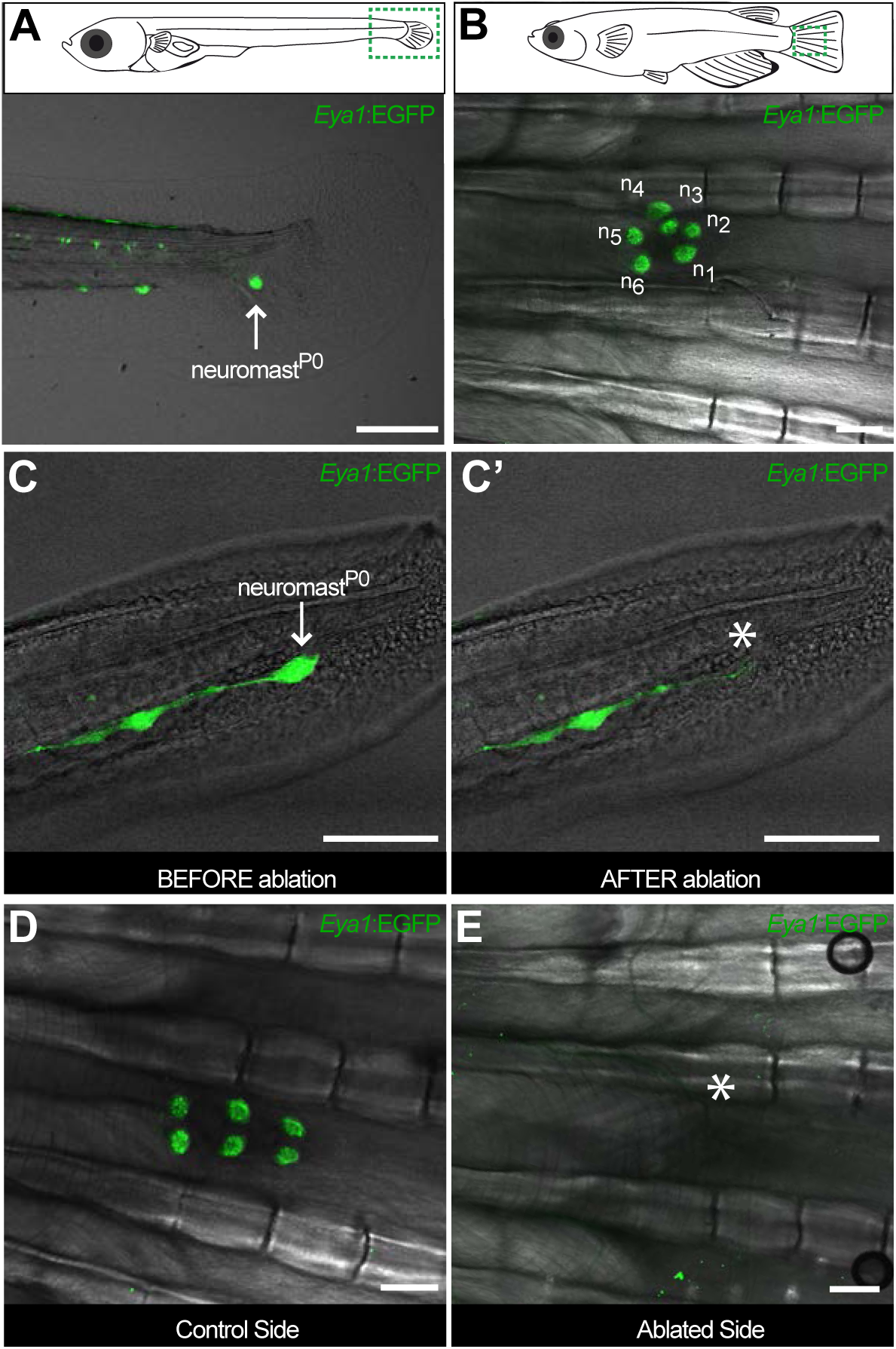
The CNC allows studying stem cells during organ growth, homeostasis and post-embryonic organ formation. Early juvenile medaka display a single neuromast on the caudal fin (neuromast^P0^, arrow in A), which gives rise to a cluster of neuromasts (n1 to n6, B) during post-embryonic life. Two-photon laser microscopy can be used to effectively ablate neuromast^P0^ (C and asterisk in C’). The caudal neuromast cluster (D) can not form in the absence of neuromast^P0^ (asterisk in E). Scalebars are 10*μ*m in embryos and 100*μ*m in adults.

We induced recombination in late embryos that were grown for two and up to 18 months post induction (Figure 5A). The analysis of long-term clones on the CNC revealed that both neural and nBC lineages are maintained by dedicated, fate-restricted stem cells (Figure 5B-D). We analysed 200 neuromasts distributed in 34 labelled, mosaic CNCs and found long-term clones in either the neural lineage (14%)(Figure 5B, D) or in the nBC lineage (24%)(Figure 5C, D). Additionally, a small proportion presented labelled cells in both nBC and neural lineages (4.6%). These co-labelled cases could be the result of simultaneous recombination of cells from both lineages or alternatively, produced by a rare bipotent stem cell. By focusing on CNCs with sparse labelling (≤ 50% of neuromasts labelled per cluster), the ratio of co-labelled clones drops further (2%, N=100), and disappears when we select for even lower recombination efficiency (≤ 25% of neuromasts labelled per cluster). The presence of fate-restricted stem cells is further supported by the existence of CNCs in which all neuromasts are labelled in either the BC (N=55 neuromasts in 12 CNCs) or the neural lineage (N=3 neuromasts in 1 CNC)(Supplementary Figure 1). Taken together, our results indicate the existence of independent lineages during neuromast homeostasis and post-embryonic organogenesis, and position the neuromast as a minimal system to tackle stem cell fate-restriction and clonal organisation in a composite organ.

**Figure 5.**
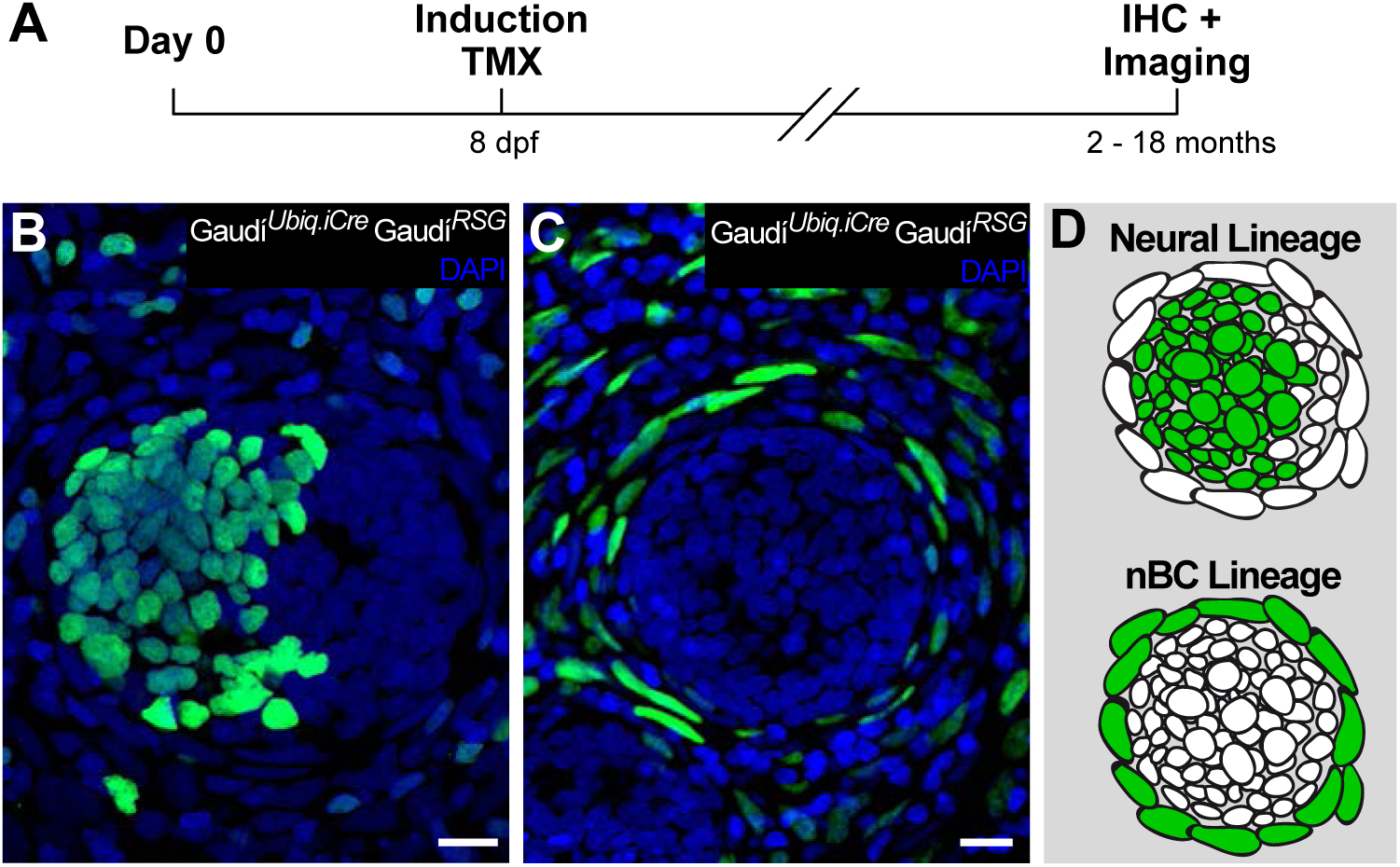
Independent stem cell populations maintain neural and nBC lineages in the mature neuromast. (A) Experimental outline of long-term lineage tracing using Gaudí^RSG^ Gaudí^Ubiq.iCre^. Neuromasts in the CNC of induced fish are labelled either in the neural lineage (B, upper scheme in D) or in the nBC lineage (C, bottom scheme in D). Neuromasts labelled in the neural lineage contain mantle, support and hair cells (B), while clones in the nBC lineage do not contribute to the neural lineage (C). Scalebars are 10*μ*m.

### Mantle cells are neural stem cells

Having shown that neuromasts contain stem cells that maintain the neural lineage, we tackled neuromast stem cell identity using a regeneration approach. Combining *Tg(K15:H2B-RFP)* with *Tg(Eya1:EGFP)* or *Tg(Eya1:H2BGFP)*, we ablated a major proportion (40% up to 95%) of neural lineage cells using two-photon laser confocal microscopy sparing a few intact *K15^+^* mantle cells (N=18 neuromasts in 4 fish)(Figure 6A-B’). Iterative post-injury imaging revealed that the surviving mantle cells initially coalesce and then proceed to re-enter the cell cycle, which results in a small circular cluster of cells (Figure 6C-C”). This regenerating neuromast progressively increases in size and eventually shows *Eya1* positive internal SCs and HCs (Figure 6D-D”). Notably, as few as four *K15^+^* cells were sufficient to regenerate the entire neural lineage of a neuromast. These results indicate that mantle cells have the potential of reconstituting all the cell types within the neural lineage of an organ after severe injury.

**Figure 6.**
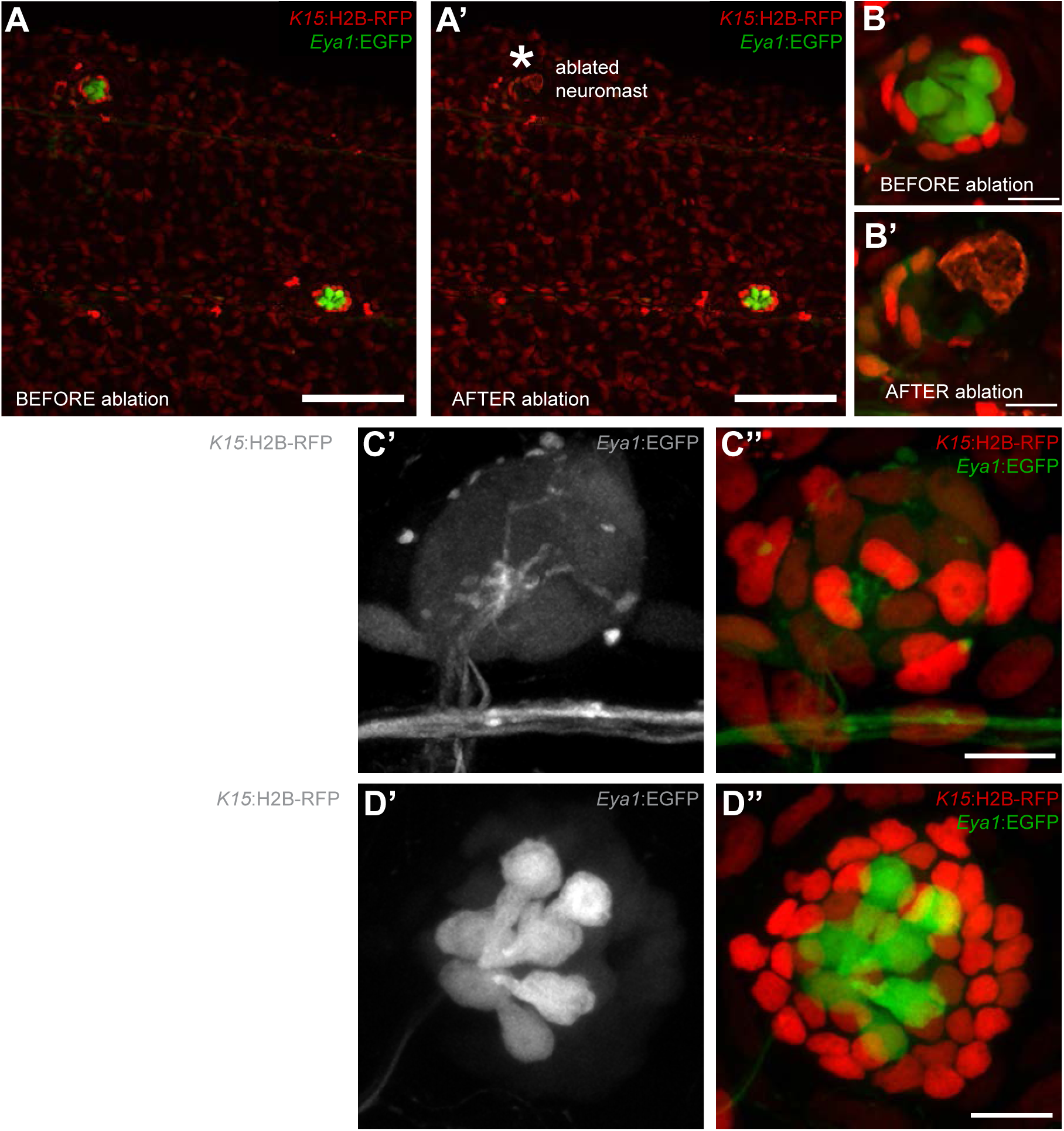
Mantle cells regenerate all cell types within the neural lineage upon severe injury. Two photon ablation on double transgenic Tg(*K15*:H2B-RFP), Tg(*Eya1*:EGFP) at 12 dpf. (A-B’). Ablations were done to remove most cells in the neuromasts, sparing a few *K15*^+^ cells (B’). The same neuromast shown 40 hours post-ablation reveals a small cluster of RFP+ cells that have coalesced around the site of injury without any apparent differentiation (C-C”). Six days post-injury all cell types within the neural lineage have been reconstituted (D-D”). Mantle and support cells can be observed in *K15*:H2B-RFP (D, D”), while hair cells and internal support cells are evident in the *Eya1*:EGFP (D’, D”). Scalebars are 100*μ*m for trunks (A, A’) and 10*μ*m for neuromasts (B-D”).

To assess whether *K15^+^* mantle cells function as neural stem cells during homeostasis and post-embryonic organogenesis, we generated Tg(*K15:*Cre*^ERT2^*) to permanently label mantle cells and their progeny. When crossed to *Gaudí^RSG^* and induced for recombination, these larvae exhibited sparse recombination in skin epithelial cells and in mantle cells as expected from the Tg(*K15:*H2BEGFP)(Figure 7A-C). We exploited the location of neuromasts to perform imaging on *K15^+^* clones at 72 hours post-induction, selecting for one-two cell clones in mantle cells (Figure 7 D,E) and annotating the recombined neuromasts. These selected fish were grown for up to one month and imaged again on the same neuromasts, and our analysis revealed that mantle cell derived clones contained support cells (N=30 neuromasts in 12 fish) (Figure 7D’) and in some cases all cell types of the neural lineage (Figure 7E’) (N=8 neuromasts in 5 fish). When mantle cell clones were detected in the neuromast^P0^ (Figure 7F), we observed that the CNC cluster formed from it contained other neuromasts with EGFP positive cells in the entire neural lineage (3 out of 5 neuromasts)(Figure 7F’,F”), indicating that *K15^+^* mantle cells also function as founder cells for post-embryonic organogenesis. Altogether, our results indicate that *K15^+^* mantle cells function as neuromast neural stem cells during homeostatic growth, post-embryonic formation of new neuromasts and organ regeneration.

**Figure 7.**
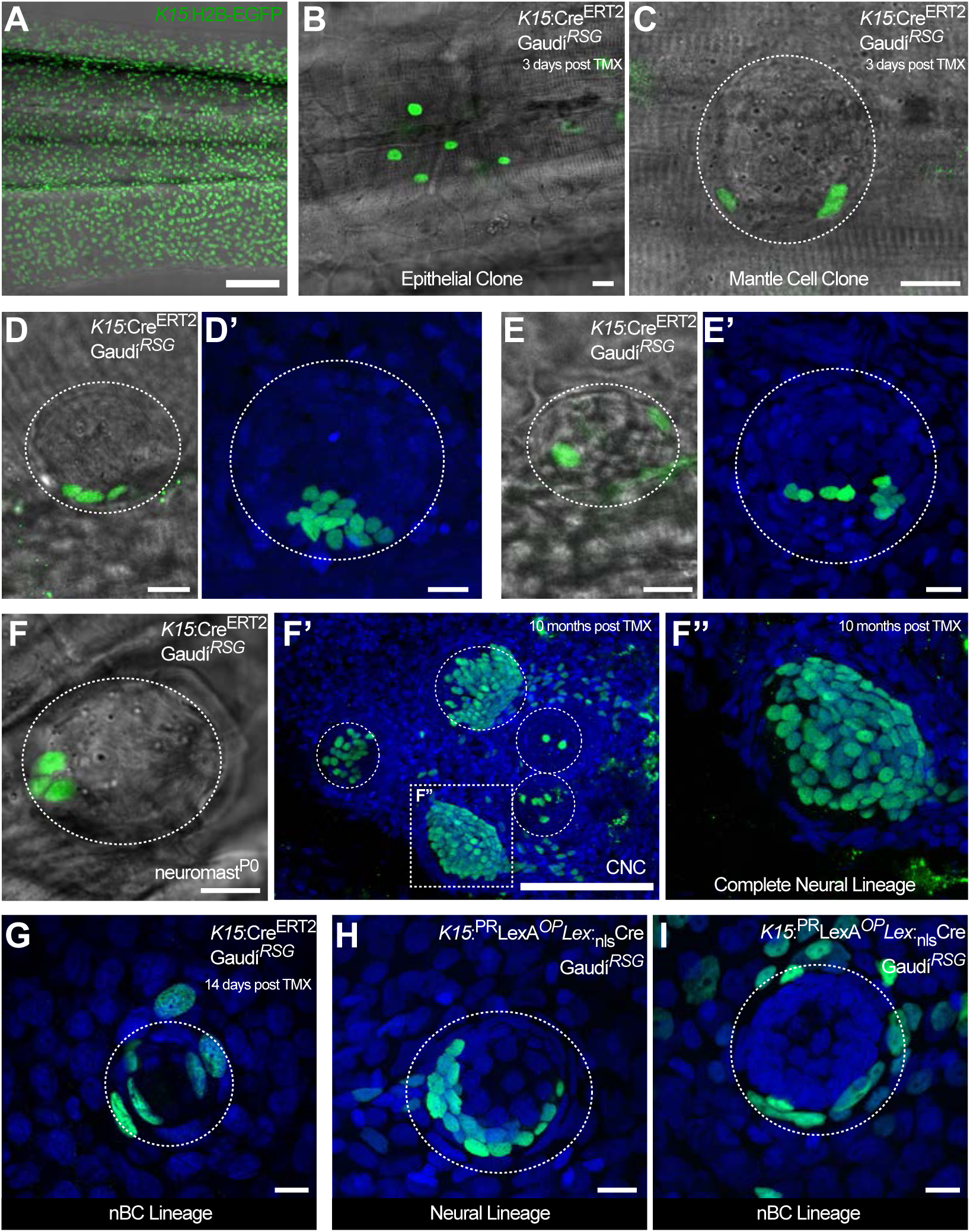
*K15^+^* mantle cells are neuromast neural stem cells. Tamoxifen induction of *K15*^+^:*^ERt2^*Cre, *Gaudí^RSG^* labels a subset of the EGFP+ cells by Tg(*K15*:H2B-EGFP) in the skin epithelium and the neuromast (Compare A to B, C). A clone of three mantle cells in induced *K15:*Cre*^ERT2^*, *Gaudí^RSG^* (D) generates more mantle and also support cells (D’). A second clone of labelled mantle cells (E) expanded into support and differentiated hair cells (E’). A clone of three mantle cells in *neuromast^P0^* (F) contributes to 3 different neuromasts in the adult caudal neuromast cluster (F’) and generate all cell types of the neural lineage (F”). A different population of *K15^+^* mantle cells generate also nBCs (G-I). Recombination of *K15:*Cre*^ERT2^*, *Gaudí^RSG^* produces some neuromasts labelled in the nBC lineage (G). Injection of a *K15*:*^PR^*LexA *^OP^*Lex:*^nls^*CRE construct into *Gaudí^RSG^* embryos generate clones either in the neural (H) or in the nBC (I) lineage. Wide scalebars are 100,pm for trunks (A, F’) and thinner scalebars are 10*μ*m.

### Independent embryonic origins for border and neural lineage

When analysing recombined clones in *Gaudí^RSG^K15:*Cre*^ERT2^* medaka that were induced during late embryogenesis, we noticed the presence of clones in the border cell lineage (Figure 7G)(N=6 neuromasts in 4 fish). This is compatible with the transient expression of EGFP we had previously observed in Tg(*K15*:H2B-EGFP) hatchlings, and suggests that nBCs indeed derive from *K15^+^* cells. The fact that the same marker labels fate-restricted stem cells for two different lineages can be explained by two possible scenarios. Neuromast BCs could originate either from a subset of *K15^+^* mantle cells that are committed to exclusively produce border cells, or from *K15^+^* cells outside of the neuromast neural lineage.

To address nBC origin, we generated mosaic embryos by injecting a *K15*:H2B-EGFP plasmid at the 2-4 cell stage into wild types. Injected embryos at 9 dpf were stained with antibodies against EGFP to allow a longer time window analysing the progeny of *K15^+^* cells. This revealed neuromasts that were either labelled in the neural lineage (N=15/37 neuromasts in 7 fish) or in the nBC lineage(N=11/37 neuromasts in 7 fish) (Supplementary Figure 2) suggesting that the two lineages are already set apart by the time of organ formation. Expectedly, we also observed neuromasts in which both border cells and the neural lineage were labelled (N=11/37 neuromasts in 7 fish) mirroring our observations with long term lineage analysis (data not shown). To follow *K15^+^* progeny for longer periods, we injected a *K15*::Lex^PR^ *Lex^OP^*:CRE plasmid into *Gaudí^RSG^* since the LexPR system provides an amplification step (Emelyanov and Parinov, 2008; Lust *et al.*, 2016) that results in higher recombination rates than the *K15*:Cre*^ERT2^* approach (see Materials and Methods). Here again, we observed neuromasts that were either labelled in the neural lineage (N=3/13 neuromasts)(Figure 7H) or in the border cell lineage (N=9/13 neuromasts)(Figure 7I). Notably, we observed that clones containing border cells reproducibly continue into skin epithelial cells in the vicinity of the neuromasts both in injected embryos (100%, N=40 neuromasts)(Figure 7I’) as well as in the previously described long term lineage analysis (100%, N>100 neuromasts)(Figure 7G, 5C). Taken together, these results indicate that nBCs could originate from *K15^+^* cells within the epithelium.

### nBCs originate from K15^+^ cells in the epithelium

To dynamically address the developmental origin of border cells, we followed a 4D approach using Tg(*K15*:H2B-EGFP) embryos. We exploited the migration of the developing midline neuromasts to their final destination (Seleit *et al.*, 2017) to follow the two *K15^+^* populations during organogenesis (Figure 8A and Supplementary Movies 2-4). As the developing midline neuromasts reach the horizontal myoseptum they come into contact with the overlying epithelial cells. Promptly, three to four *K15^+^* epithelial cells (red, green and blue dots in Figure 8A) respond to the arrival of the neural stem cell precursors. Over a period of 72 hours, these epithelial cells undergo significant morphological changes resulting in the characteristic elongated shape of border cell nuclei. Both ventral and midline neuromasts of the posterior lateral line system in medaka, as well as neuromasts of the anterior lateral line (aLL), trigger the same behaviour in the surrounding epithelium (5 ventral neuromasts and 4 midline neuromasts in the pLL and 2 neuromasts in the aLL in 5 embryos). We wondered whether in addition to a change in nuclear shape, the cellular membranes of transforming cells also undergo morphological remodelling. We performed double injections of *K15*:H2B-RFP and *K1 5*:mYFP plasmids into *Cab* wild type embryos to create clones labelled for nuclei and membranes. The results reveal that a change in cellular morphology accompanies a change in nuclear shape (Figure 8B, B’, white and green asterisks) (quantifications in Supplementary Figure 3). Taken together, these results prove that nBCs originate from transformed *K15^+^* skin epithelial cells.

**Figure 8.**
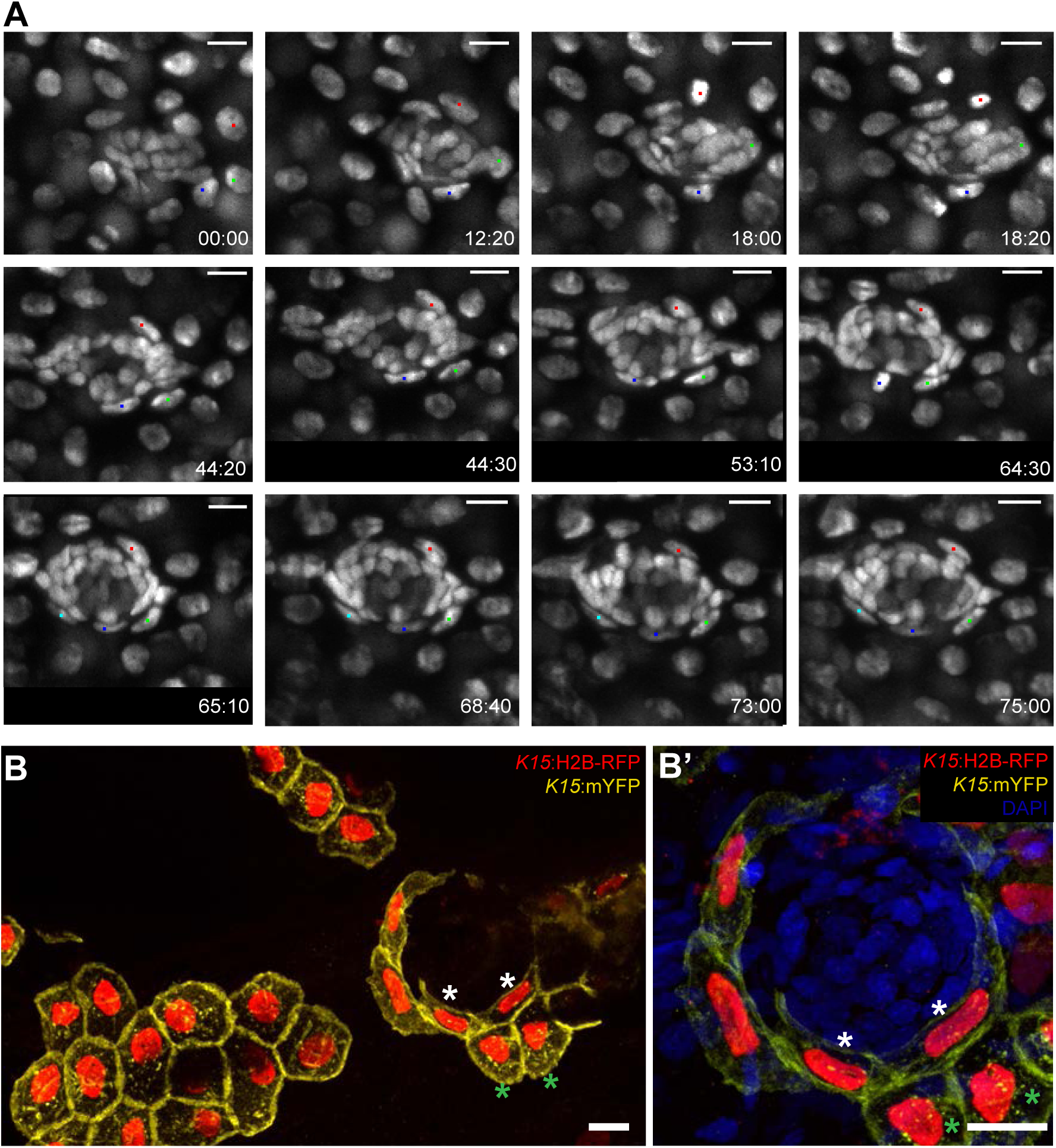
Transformation of skin epithelial cells into nBCs during neuromast formation. (A) Time-lapse imaging of a stage 35 Tg(*K15*:H2B-EGFP) embryo during secondary organ formation, where three epithelial cells (blue, green and red dots) are seen to dynamically associate with the forming neuromast. The red cell divides (18:00h to 18:20h) before the morphological transformation, generating one skin epithelial daughter and another daughter that will transform into a nBC (64:00h to 75:00h). The green cell transforms into a nBC without diving, and the blue cell first transforms (18:20h to 43:00h) and then divides to generate two nBC (64:30h to 65:10h) that stay in the neuromast. The images in (A) are selected time-points from three consecutive movies (see Supplementary Movies) of the same developing neuromast. (B-B’) Immunostaining of a double injected (mosaic) *K15*:mYFP, *K15*:H2B-RFP embryo shows that nBCs transformation involves a drastic remodelling of both nuclei and cellular morphology (B). Compare skin epithelial cells (green asterisks in B, B’) with their sibling nBCs (white asterisks in B,B’) included in the neuromast. Scalebars are 10*μ*m.

### The neural lineage is necessary and sufficient for epithelial transformation

Neuromast organogenesis requires that cells deposited by the primordium (neuromast neural lineage) and skin epithelial cells come into contact. This could happen either by an active migration of cells deposited by the primordium towards pre-defined hot spots for epithelial transformation or alternatively, by the local transformation of epithelial cells induced by the unspecified arrival of the neuromast neural lineage. Our previous observations suggest a leading role for the neural lineage, since left and right pLLs in medaka larvae often have an unequal number of neuromasts located in undetermined positions (Seleit *et al.*, 2017). Irrespective of how many neural clusters are generated by the primordium, and regardless of the final position along the anterior-posterior axis, all mature neuromasts contain nBCs.

We experimentally tackled the hierarchical organisation of the interacting tissues by two complementary approaches that result in fish with either ectopic or absent neuromasts along the pLL. The loss-of-function of *neurogenin* or *erbb* has been shown to result in the formation of ectopic neuromasts in zebrafish by neural lineage interneuromast cells that re-enter the cell cycle (Grant *et al.*, 2005; López-Schier and Hudspeth, 2005; Lush and Piotrowski, 2014). The injection of Cas9 mRNA and two gRNAs directed against medaka *neurogenin* into Tg(*Eya1*:EGFP) or Tg(*Eya1*:H2B-EGFP) resulted in juveniles with an increased amount of pLL neuromasts (Figure 9A). We focused on the severest phenotypes, which contained twice the amount of neuromasts as compared to their control siblings (N=58 pLL neuromasts in 2 larvae). Our results show that ectopic midline neuromasts are composed of neural lineage and nBCs (Figure 9B-H) (N=33 neuromasts in 2 larvae). This suggests that the presence of the neural lineage is sufficient to drive ectopic transformation of epithelial cells into border cells. Complementarily, ablation of the primordium before deposition of the last neuromast results in fish lacking the CNC (Figure 9I,J). The stereotypic position of the CNC constitutes an ideal set up to explore whether transformation of epithelial cells into border cells occurs even in the absence of the neural lineage. A detailed analysis of DAPI stained caudal fins revealed the absence of transformed border cells on the ablated side, as opposed to their presence on the non-ablated, control side (Figure 9I’,J’). Altogether, our results demonstrate that the neural lineage is both necessary and sufficient to induce the transformation of epithelial cells into neuromast border cells *in situ*.

**Figure 9.**
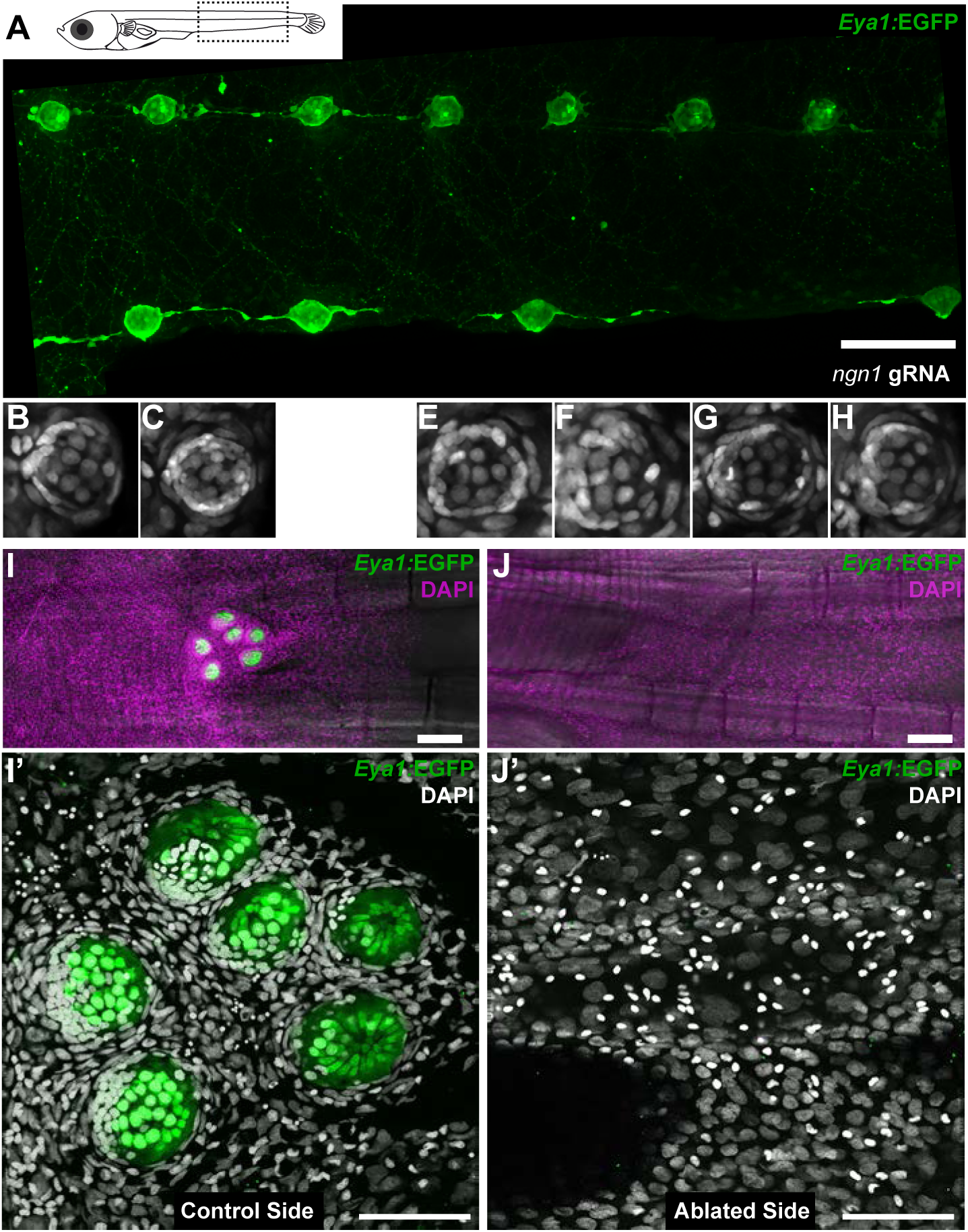
Arrival of neural neuromast cells is necessary and sufficient to induce transformation of epithelial cells into nBCs. (A) Anti-GFP staining of a Tg(*Eya1*:EGFP) embryo at stage 36 that was injected with *ngn1* gRNAs reveals the formation of ectopic pLL midline neuromasts. (B-H) DAPI images of seven consecutive midline neuromasts show that all of them contain nBCs. (I-J’) Absence of neural lineage prevents the transformation of the skin epithelium. While each neuromast of a control CNC (I) contains nBCs (I’), early ablation of neural lineage in neuromast^P0^ results in no CNC (J) and no skin epithelium transformation (J’). Wide scalebars are 100*μ*m (A, I) and thinner scalebars are 50*μ*m (I’, J’).

## Discussion

Here we identify a new type of neural stem cell (*Keratin15*^+^) that fulfils all criteria of stemness: it generates more neural stem cells, all neuromast neural progenitors and neurons during homeostasis and regeneration and additionally, functions as the organ founder cell during post-embryonic organogenesis. We tracked these NSCs back to organogenesis to show that their precursors modify their environment to transform and recruit a new cell type to the forming organ.

### The origin of lineage commitment

Lineage commitment in stem cells could result from alternative scenarios. On the one hand, stem cells with different potencies can derive from a common pool of parental cells that acquire fate-restriction as development proceeds. This is the case for luminal (*K8^+^*) and myoepithelial (*K14^+^*) fate-restricted stem cells in the mammary gland that derive from a common pool of (*K14^+^*) embryonic multipotent cells (Van Keymeulen *et al.*, 2011), and neural retina and pigmented epithelium (*Rx2^+^*) fate-restricted stem cells in the post-embryonic fish retina that derive from common embryonic retinal progenitors cells (Centanin *et al.*, 2014; 2011; Reinhardt *et al.*, 2015). On the other hand, stem cells with independent embryonic origins could be brought together during early organogenesis to maintain different tissues in the same organ, as typically observed in the mammalian skin (Fuchs *et al.*, 2004; Ouspenskaia *et al.*, 2016) and more generally, in most composite organs. We have presented evidence that indicates a dual embryonic origin for medaka neuromasts. A neuromast neural lineage consists of mantle, support and hair cells surrounded by a second, independent lineage of neuromast border cells. Our findings position neuromasts as an approachable and minimalist system to address the coordination of independent lineages during homeostatic replacement, regeneration and post-embryonic organogenesis.

### Mantle cells during regeneration

We observed that a few remaining *K15^+^* mantle cells can reconstitute the entire neural lineage upon severe injury. Interestingly, the dynamics of neuromast regeneration seems to resemble the formation of secondary neuromasts during embryonic development (Seleit *et al.*, 2017). We observe that upon injury the initial reaction of the surviving mantle cells is not to proliferate but rather to coalesce, followed by an amplification of cell numbers. Only afterwards does differentiation take place and all three cell types of the neural lineage are reconstituted. This highly ordered transition suggests that before differentiation can take place developing neuromasts must attain a critical number of cells, as we observed during embryonic development (Seleit *et al.*, 2017). Intriguingly, the efficient regeneration of neuromasts in medaka contrasts with its reported inability to adequately regenerate its heart upon mechanical injury (Ito *et al.*, 2014). It has recently been shown that regeneration enhancer elements exist in zebrafish and can drive tissue specific regenerative responses to injury (Kang *et al.*, 2016). A differential selective pressure could impact those TREEs (tissue-specific regeneration enhancer elements) in a species-specific manner, and can therefore have divergent effects on the regenerative capacity of specific organs. A comparative genomic approach on TREEs among teleosts could contribute to the understanding of their differential regenerative capacities, and our results provide yet another module to study the evolution of regeneration responses across fish.

### Mantle cells as neuromasts stem cells

Neuromasts in fish can replace lost HCs during their entire life, which has been taken as an indication that neuromasts should contain neural stem cells. Numerous short-term studies in zebrafish, however, have reported that both during homeostasis and regeneration HCs are produced by post-mitotic hair cells precursors (Hernández *et al.*, 2007; Kniss *et al.*, 2016) and proliferation of support cells (Hernández *et al.*, 2007; López-Schier and Hudspeth, 2006; Pinto-Teixeira *et al.*, 2015; Romero-Carvajal *et al.*, 2015). In the past years, several studies have reported the existence of compartments or territories within the neuromasts that permit either self-renewal of support cells or differentiation into HCs (Pinto-Teixeira *et al.*, 2015; Romero-Carvajal *et al.*, 2015; Wibowo *et al.*, 2011). Despite the detailed knowledge on short-term aspects of regeneration and homeostatic replacement, the existence and identity of long-term neuromast stem cells has remained elusive. Using genetic tools to irreversibly label cells and their progeny, we followed long-term lineages to show the existence of neural stem cells in the organ and uncover their identity. Using ubiquitous drivers for recombination (either *Ubiquitin*:Cre*^ERT2^* or *Hsp70*:_nls_Cre) we observed that all long-term clones in the neural lineage contain mantle cells, suggesting that these are necessary for the clone to be maintained. Additionally, a cell-type specific driver for mantle cells (*K15:*Cre*^ERT2^*) proved these cells to function as neural stem cells under homeostatic conditions. Our results indicate that *K15^+^* mantle cells in medaka work as *bona fide* neural stem cells, maintaining clones that expand within neuromasts and generating new organs during post-embryonic life.

### Induced fate transformation

We report that upon arrival of neuromast neural lineage cells deposited by the migrating primordium (the case of ventral neuromasts) or originated by coalescence of inter-neuromast cells (the case of midline neuromasts), epithelial cells follow a morphological and molecular transformation that results in the generation of border cells. The transformation event seems to be restricted to the initial phases of organ formation and involves a small number of cells. Our 4D analysis of the epithelial-to-border cell transformation has revealed that border cells can divide once transformed to generate two border cells, which was confirmed by short IdU pulses (E. Ambrosio, *not shown*). Labelled cells can generate large clones that involve all border cells in every neuromast of the CNC, indicating a long-term stability of labelled clones. Altogether, we have shown that neural stem cell precursors trigger the transformation of epithelial cells, which will in turn associate with them to form a mature neuromast organ. We believe that our findings position neuromasts as a new paradigm to dynamically study the inductive behaviour that cell types exert on each other during organogenesis.

The formation of the lens in the vertebrate retina was the first reported case of tissue induction. There, the developing optic vesicle (composed of neural progenitors) contacts the surface ectoderm to induce a lens placode from previous ectoderm cells. The case that we report here shares the same rationale and interestingly, the cells involved express the same molecular markers. Neuromast neural lineage cells are *Eya1*^+^ as retinal progenitors, and they induce the transformation of *K15*^+^ ectodermal cells. The similar molecular identity of the cell types involved, in which *Eya1^+^* cells operate as the inducer and *K15*^+^*Eya1*^-^ cells are the responder, invites to question whether similar molecular mechanisms are involved in the transformation event. The identity and nature of the inducer cells — in both cases neural progenitors responsible for generating neurons that define the organ’s function — indicate a shared hierarchical organisation during organogenesis of sensory systems. It would be interesting to address whether other sensory systems with a similar surface location and common molecular markers (e.g. the ampular system) follow the same rationale during organogenesis and ultimately, whether proximate interaction, or induction, constitutes an ancient signature for the establishment of sensory systems.

### Do nBCs constitute a niche for neural stem cells?

The physical proximity of neuromast border cells and mantle cells, and the direct protrusions between them observed in our EM images raise the possibility that nBCs constitute a niche for the neural stem cells of the organ. The concept of the stem cell niche was introduced by Schofield during the late 70’s (Schofield, 1978). The niche was proposed to influence the behaviour of stem cells by a variety of mechanisms including direct signalling, in a way that stem cells that kept a close connection to the niche were destined to maintain their stemness while their displaced progeny start to differentiate. Indeed, we observe that MCs maintaining contact with nBCs keep expression of a neural stem cell marker (*K15^+^*) and display stem cell behaviour, i.e. generation of long-term clones. Cells located to the interior of the organ, and therefore away from the outermost nBCs, progressively lose expression of the stem cell marker to eventually acquire molecular and morphological features of differentiating neurons.

The existence of niches was reported in the majority of stem cell systems, ranging from plants to animals and from invertebrates to vertebrates. Most niche cells have a different lineage than their respective stem cells (Fuchs *et al.*, 2004), which raises fundamental developmental questions like how do these cell types come together (Tamplin *et al.*, 2015), whether they are formed simultaneously or sequentially (Ouspenskaia *et al.*, 2016) and if there is a hierarchy organising their interaction. Our results suggest that during development, the neural lineage induces the formation of its own niche by fate conversion of neighbouring epithelial cells. The induction of a transient, short-lived niche by HSCs in zebrafish was recently reported to occur by a major morphological remodelling of perivascular endothelial cells (Tamplin *et al.*, 2015). We report a similar morphological transformation during sensory organ formation that notably results in a life-long, permanent niche. This model in which niche formation is triggered by arriving neural precursors seems appropriate for a system in which the number and location of organs is plastic and not defined genetically. Whether the same rationale applies to niche formation upon the arrival of migrating malignant cells in pathological cases is a clinically relevant question that remains to be elucidated.

## Material and Methods

### Fish stocks and generation of transgenic lines

Medaka (*Oryzias latipes*) stocks were maintained as closed stocks in a fish facility built according to the local animal welfare standards (Tierschutzgesetz § 11, Abs. 1, Nr. 1), and animal experiments were performed in accordance with European Union animal welfare guidelines. The facility is under the supervision of the local representative of the animal welfare agency. Fish were maintained in a constant recirculating system at 28°C with a 14 h light/10 h dark cycle (Tierschutzgesetz 111, Abs. 1, Nr. 1, Haltungserlaubnis AZ35–9185.64 and AZ35–9185.64/BH KIT).

The strains and transgenic lines used in this study are: Cab (medaka Southern wild type population), Tg(*Eya1*:EGFP) (Seleit *et al.*, 2017), Gaudí^Ubiq.iCre^, Gaudí^Hsp70.A^, Gaudí^RSG^ (Centanin *et al.*, 2014). The following transgenic lines were generated for this manuscript by I-SceI mediated insertion, as previously described (Rembold *et al.*, 2006): Tg(*K15*:mYFP), Tg(*K15*:H2B-EGFP), Tg(*K15*:H2BRFP), Tg(*K15*:Cre*^ERT2^*), Tg(*neurom^K8^*:H2B-EGFP).

#### Generation of the constructs *K15*:mYFP, *K15*:H2B-EGFP, *K15*:H2B-RFP

A 2.3 kb fragment upstream of the medaka *Keratin15* ATG was amplified by PCR using specific primers flanked by XhoI sites (forward: ACTGACTCGAGACCAAAGGAAAGCAGATGAA; reverse: ACTGACTCGAGTTGTGCAGTGTGGTCGGAGA) using the fosmid GOLWFno691_n05 (NBRP Medaka) as template. The PCR fragment was cloned into an I-SceI vector already containing mYFP, and sub-cloned from there into I-SceI vectors containing either H2B-EGFP or H2B-RFP.

#### Generation of the construct *K15:*Cre*^ERT2^*

The 2.3Kb *Keratin15* promoter was cut with XhoI from the *K15*:mYFP plasmid and cloned upstream of Cre*^ERT2^*, replacing the ubiquitin promotor in *Ubiquitin*:Cre*^ERT2^* (Centanin *et al.*, 2014).

#### Generation of the construct *K8*:H2B-EGFP

The 0.5Kb *Keratin8* promoter from zebrafish (Emelyanov and Parinov, 2008) was sub-cloned into an I-SceI vector containing H2B-EGFP via KpnI/AscI. Among the founders obtained, one expressed high levels of H2B-EGFP in both skin epithelium and neuromasts and was used to establish Tg(*neurom^K8^*:H2B-EGFP). Other founders from the same injection did not share the expression in the neuromasts.

### Generation of clones

Fish from the Gaudí^RSG^ transgenic line were crossed with either Gaudí^Ubiq.iCre^ or Tg(K15:Cre*^ERT2^*). The progeny from these crosses was induced with a 5µM tamoxifen (T5648 Sigma) solution for 3-12h and afterwards washed extensively with ERM. When Gaudí^RSG^ fish were crossed to Gaudí^Hsp70.A^, double transgenic embryos were heat-shocked using ERM at 42°C and transferred to 37°C for 1-3h.

Recombined fish were selected under a fluorescent scope and imaged afterwards. For short-term analysis, t1 was 3-8 and t2 15-27 dpf. For long-term analysis, fish were grown for additional 2-18 months post induction. Cell type annotation was done based on nuclear morphology and position within the neuromast. In long-term lineage analysis, we quantified clones bigger than 4 cells. Clones generated by injection of DNA into the 1 - 2 cell stage were prepared as previously stated (Rembold *et al.*, 2006).

### Antibodies and staining

Immunofluorescence staining were performed as previously described (Centanin *et al.*, 2014). The primary antibodies used in this study were rabbit a-GFP, chicken a-GFP (Invitrogen, both 1/750) and rabbit a-Sox2 (GeneTex, 1/100). Secondary antibodies were Alexa 488 a-Rabbit, Alexa 546 a-Rabbit, Alexa 647 a-Rabbit (Invitrogen, all 1/500). DAPI was used in a final concentration of 5ug/l.

### Imaging and image analysis

#### Preparation of samples for live imaging

Embryos were prepared for live imaging as previously described (Seleit *et al.*, 2017). Briefly, we used a 20 mg/ml as a 20x stock solution of tricaine (Sigma-Aldrich, A5040-25G) to anaesthetise dechorionated embryos and mounted them in low melting agarose (0,6 to 1%). Imaging was done on glass-bottomed dishes (MatTek corporation).

#### Preparation of fixed samples

Stained samples were mounted in glycerol 80% on glass slides. Samples that required imaging from both sides of the fish were mounted between two cover slides using a minimal spacer.

#### Imaging

Anaesthetised embryos were screened using an Olympus MVX10 binocular coupled to a Leica DFC500 camera. For the acquisition of high quality images, we used a Nikon AZ100 scope coupled to a Nikon C1 confocal, or the laser-scanning confocal microscopes Leica TCS SPE and Leica TCS SP5 II. When imaging living samples over long-term time lapse, a Microscope Slide Temperature Controller (Biotronix) was used. Time-lapse imaging of transforming epithelial cells was done over a period of 72 hours using an EMBL MuVi-SPIM (Krzic *et al.*, 2012) with two illumination objectives (10x Nikon Plan Fluorite Objective, 0.30 NA) and two detection objectives (16X Nikon CFI LWD Plan Fluorite Objective, 0.80 NA). Embryos were placed in glass capillaries using 0,6% low melting agarose. All subsequent image analysis was performed using standard Fiji software.

#### Image analysis

We used the free standard Fiji software for the analysis and edition of most images. Stitching was performed automatically using 2D and 3D stitching plug-ins on ImageJ or using Adobe Photoshop to align images manually.

### EM

10dpf. Tg(*Eya1*:EGFP) embryos were fixed in 2.5% glutaraldehyde and 4% paraformaldehyde in 0.1M PHEM buffer for 30 min at room temperature and at 4°C overnight. After rinsing in buffer, embryos were imaged under a Leica MZ 10F stereo microscope for fluorescent imaging (Leica Microsystems, Vienna) to localise neuromasts. The samples were further fixed in 1% osmium in 0.1M PHEM buffer, washed in water, and incubated in 1% uranylacetate in water overnight. Dehydration was done in 10 min steps in an acetone gradient followed by stepwise Spurr resin infiltration at room temperature and polymerization at 60°C. The resulting blocks were trimmed around the neuromast to get longitudinal sections of the nBCs surrounding the mantle cells and sectioned using a leica UC6 ultramicrotome (Leica Microsystems Vienna) in 70nm thin sections. The sections were placed on formvar coated slot grids, post-stained and imaged on a JEOL JEM1400 electron microscope (JEOL, Tokyo) operating at 80 kV and equipped with a 4K TemCam F416 (Tietz Video and Image Processing Systems GmBH, Gautig).

### 2-Photon Laser Ablations

#### Ablation of neuromast^P0^

We used a multi-photon laser coupled to a Leica TCS SP5 II microscope to perform specific ablations of the neuromast^P0^ on Tg(*Eya1*:EGFP) embryos. We choose the option “Area ablations” and use the 880 nm wavelength with a laser power ranging from 25 to 30%. The absence of the neuromast^P0^ was checked immediately after the ablation and confirmed 24h later.

#### Partial ablation of neuromasts

We used a TriM Scope 2-photon microscope (LaVision BioTec, Bielefeld, Germany) as previously described (Seleit *et al.*, 2017). The 2-photon module was mounted on a Nikon FN-1 upright microscope combined with a Chameleon Ultra II femtosecond Ti:Sa laser (Coherent, Dieburg, Germany). Ablations were preformed on late Tg(*K15*:H2B-RFP) Tg(*Eya1*:EGFP) double transgenic embryos, using a 740 nm wavelength. Laser power ranged from 150 to 700 mW and was adjusted depending on the position of target neuromasts and the scope of the injury.

### Mosaic loss-of-function of ngn1

We used the Ultracontig 115 (NBRP Medaka) to design gRNAs targeting the coding sequence of medaka *ngn1*. Two gRNAs were selected (ngn1-1: UUCUCAGUGCUCGAGUCCGGCGG, and ngn1-2: UUCUCAGUGCUCGAGUCCGGCGG) using the freely available CCTop (Stemmer *et al.*, 2015). gRNA synthesis was done as previously reported (Stemmer *et al.*, 2015) using the following oligos:

gRNAngn1-1F: TAGGTTCTCAGTGCTCGAGTCCGG

gRNAngn1-1R: AAACCCGGACTCGAGCACTGAGAA

sgRNAngn1-2F: TAGGTCTGCGATGCGGATGGTCT

sgRNAngn1-2R: AAACAGACCATCCGCATCGCAGA

Tg(*Eya1*:EGFP) or Tg(*Eya1*:H2B-EGFP) embryos were injected at the 1 cell stage with a solution containing 15 ng/μl of each gRNA and 150 ng/μl of CAS9 mRNA. The resulting embryos were selected for the presence of ectopic neuromasts at late embryonic stages.

### Quantifying morphological changes

Circularity was used to measure the transformation of shape observed between epithelial cells and nBCs. Circularity = 4π(area/perimeter^2), a circularity value of 1.0 denotes a perfect circle. As the shape deforms away from a circle it’s circularity value decreases. Standard Fiji software was used to calculate the circularity of 20 nuclei and cellular membranes of epithelial cells and 20 nuclei and cellular membranes of nBCs obtained from mosaic clones of fish co-injected with the *K15*:H2BRFP and *K15*:mYFP plasmids. The generation of boxplots was done using standard R software.

## Competing interests

The authors declare no competing or financial interests.

## Acknowledgements

We thank T. Piotrowski, S. Lemke, J. Wittbrodt and M. Allende for scientific inputs on earlier versions of this manuscript, and. K. Lust, A. Gutierrez-Triana and Centanin lab members for active discussions on the project. We are grateful to J.Mateo Cerdan for help on the identification of the *Keratin 15* promoter, NBRP Medaka for sharing the fosmid containing *Keratin15* and C. Funaya, S. Gold and S. Hillmer from the EMCF facility at Heidelberg University for great technical assistance in the preparation and processing of electron microscopy data. We thank R. Bump for the analysis of clones on the CNCs, and E. Leist, A.Sarraceno and M.Majewski for assistance regarding fish maintenance. This work has been funded by the Deutsche Forschungsgemeinschaft (German Research Foundation, DFG) via the Collaborative Research Centre SFB873 (subproject A11 to L.C.) and the Cluster of Excellence Cellular Networks (Cell Networks) (to N.D.).

## Figure Legends

**Supplementary Figure 1.**
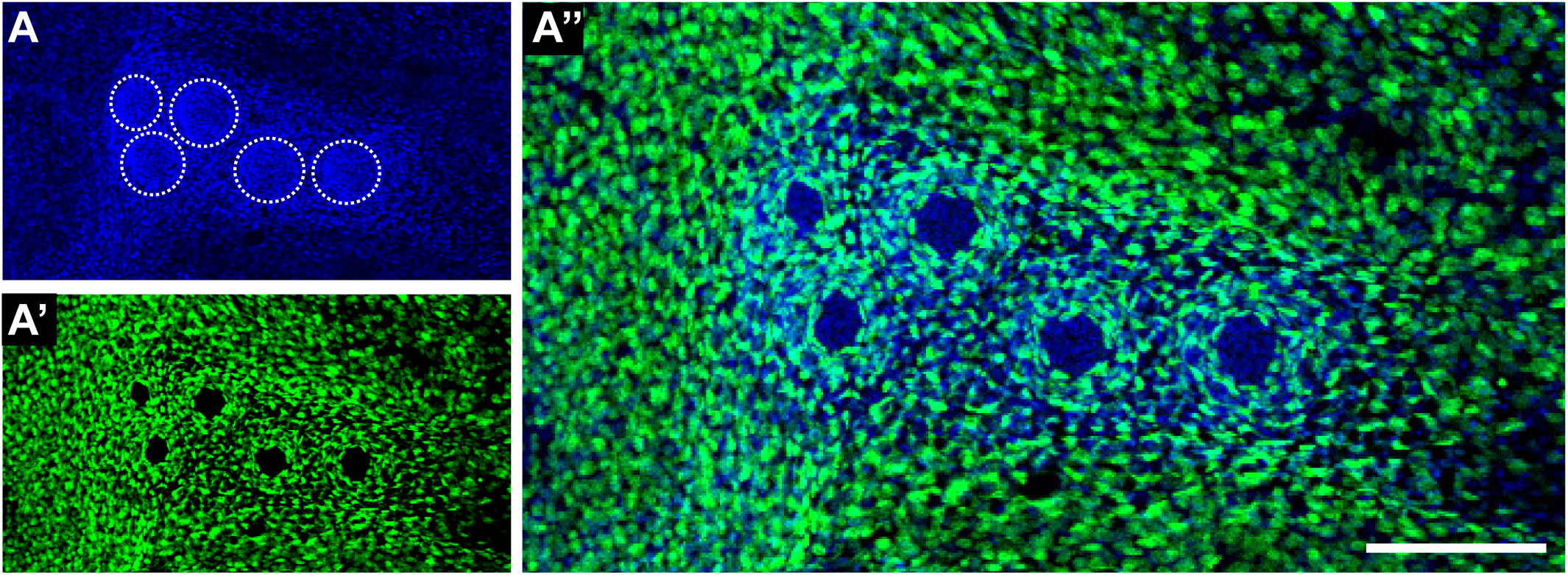
Independent lineages occur even in highly recombined CNCs. A CNC containing five neuromasts (A) shows massive recombination (A’) due to a strong induction of the double transgenic Gaudí*^RSG^* Gaudí^*Ubiq.iCre*^. Recombined cells are distributed all over the caudal fin and label nBCs in every neuromast but no cells from the neural lineage, indicating their independent origin. Scalebar is 100µm.

**Supplementary Figure 2.**
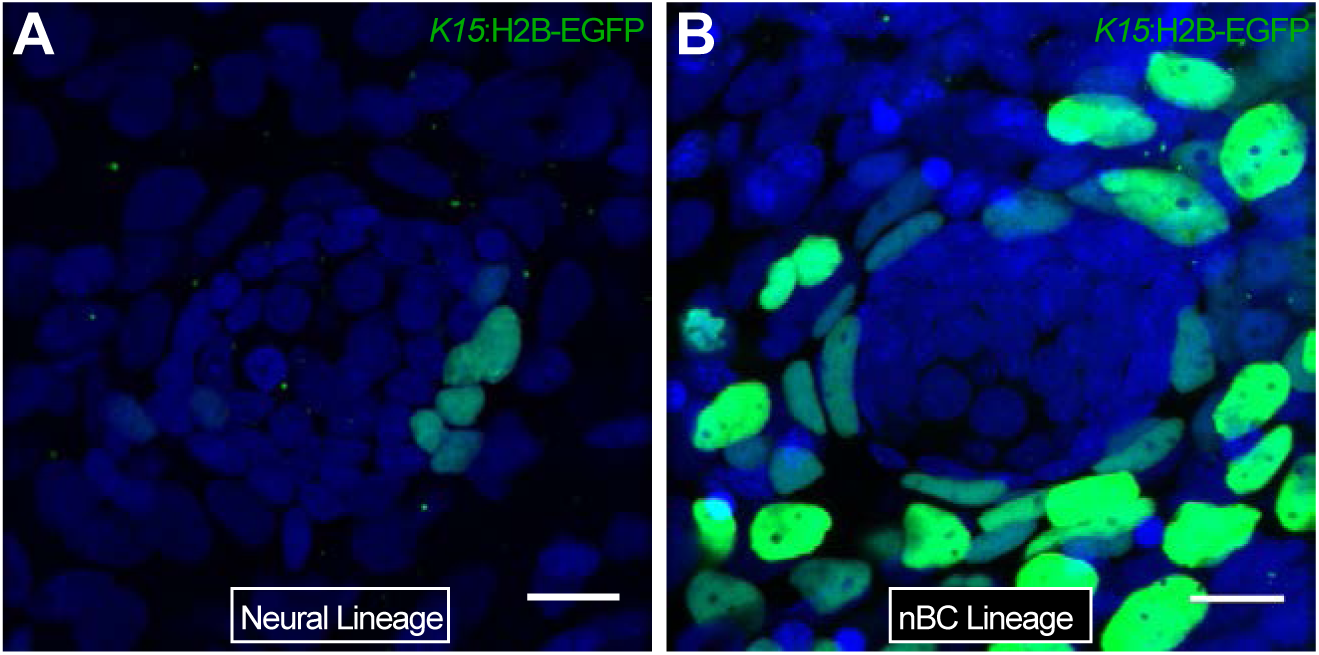
Independent origin of neural and nBC lineages in medaka neuromasts. Mosaics were generated by injection of the plasmid *K15*:H2B-EGFP at the 1-2 cell stage. Resulting clones labelled either the neural lineage (A) or the nBC lineage (B). Clones in the nBC lineage were always continious with the skin epithelium. Scale bars are 10µm.

**Supplementary Figure 3.**
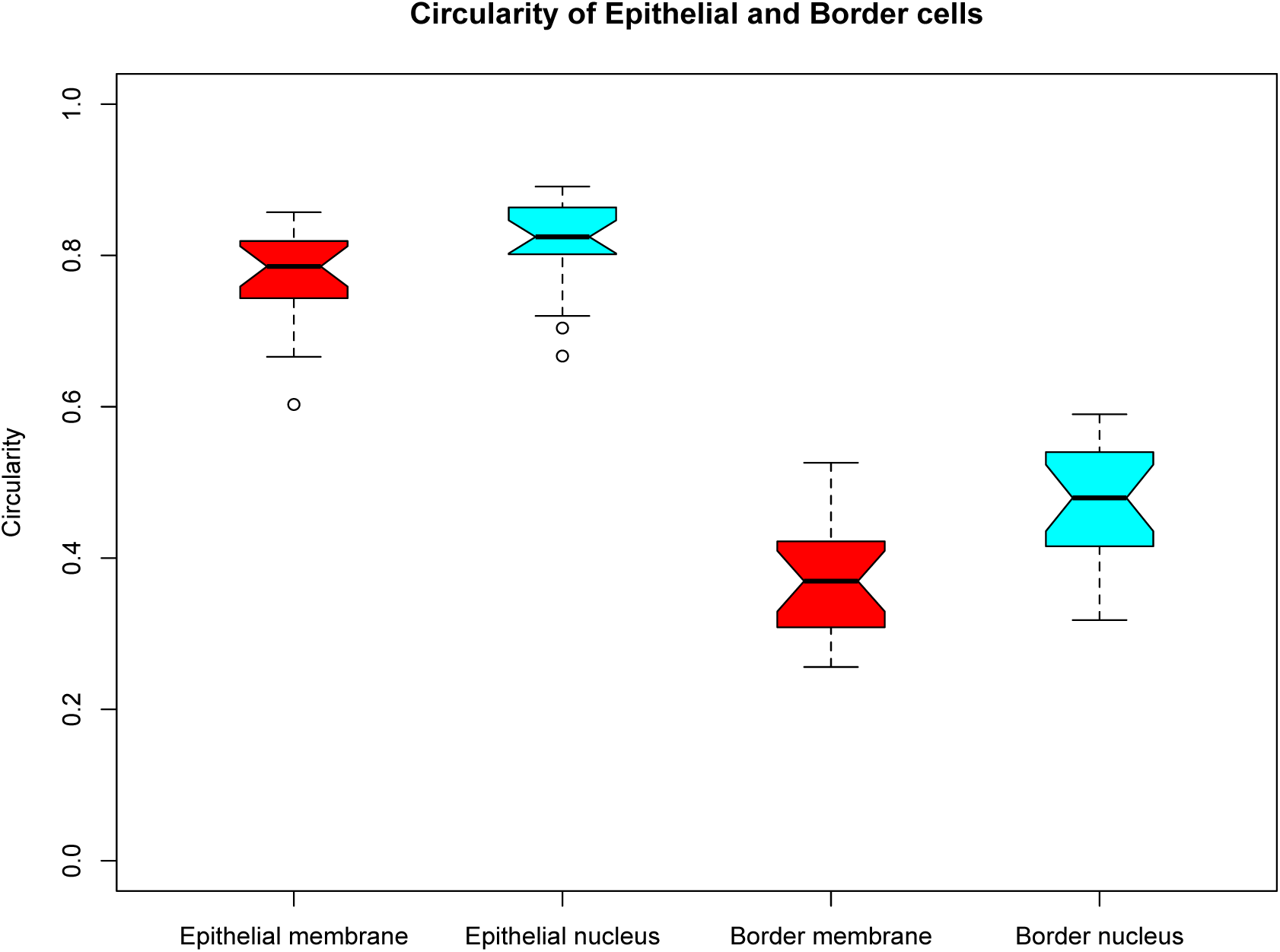
Changes in shape of cell nuclei correlate with changes in cell morphology. A circularity index was used to quantify changes in cell morpholgy and compared to that of nuclei. Cells in the skin epithelium display a more rounded morphology with more rounded nuclei than transformed nBCs.

